# Endogenous auxin directs development of embryonic stem cells into somatic proembryos in Arabidopsis

**DOI:** 10.1101/2021.08.06.455432

**Authors:** Omid Karami, Cheryl Philipsen, Arezoo Rahimi, Annisa Ratna Nurillah, Kim Boutilier, Remko Offringa

## Abstract

Somatic embryogenesis (SE) is the process by which embryos develop from *in vitro* cultured vegetative tissue explants. The synthetic auxin 2,4-dichlorophenoxyacetic acid (2,4-D) is widely used for SE induction, but SE can also be induced by overexpression of specific transcription factors, such as AT-HOOK MOTIF NUCLEAR LOCALIZED 15 (AHL15). 2,4-D and AHL15 both trigger the biosynthesis of the natural auxin indole-3-acetic acid (IAA). However, the role of this endogenously produced auxin in SE is yet not well understood. In this study we show that the induction of embryonic stem cells from explants does not require IAA biosynthesis, whereas an increase in IAA levels is essential to maintain embryo identity and for embryo formation from these stem cells. Further analysis showed that *YUCCA* (*YUC*) genes involved in the IPyA auxin biosynthesis pathway are up-regulated in embryo-forming tissues. Chemical inhibition of the IPyA pathway significantly reduced or completely inhibited the formation of somatic embryos in both 2,4-D-and AHL15-dependent systems. In the latter system, SE could be restored by exogenous IAA application, confirming that the biosynthesis-mediated increase in IAA levels is important. Our analyses also showed that PIN1 and AUX1 are the major auxin carriers that determine respectively auxin efflux and influx during SE. This auxin transport machinery is required for the proper transition of embryonic cells to proembryos and, later, for correct cell fate specification and differentiation. Taken together, our results indicate that auxin biosynthesis in conjunction with its polar transport are required during SE for multicellular somatic proembryo development and differentiation.

**One sentence summary:** Somatic embryogenesis in Arabidopsis requires auxin biosynthesis and polar auxin transport only after the acquisition of embryonic competence for somatic proembryo development and differentiation.

## Introduction

Plant growth and development is controlled to a large extent by plant growth regulators. The natural auxin, indole-3-acetic acid (IAA), is a major plant growth regulator, as it is involved in a wide array of physiological and developmental processes. An important factor determining the developmental role of auxin is its intracellular level, which is regulated by *de novo* biosynthesis, metabolism and transport. Together these three processes generate patterns of auxin maxima and minima in a cell type-dependent manner (Vanneste and Friml, 2009; Paque and Weijers, 2016).

The amino acid tryptophan (Trp) is the main precursor for IAA biosynthesis in plants. Recent genetic studies have uncovered several Trp-dependent IAA biosynthesis pathways (Tivendale et al., 2014; Zhao, 2018). Of these, the IPyA pathway has been well-characterized. The IPyA pathway consists of a two-step reaction. First, Trp is converted into indole-3-pyruvic acid (IPyA) by the TRYPTOPHAN AMINOTRANSFERASE OF ARABIDOPSIS1/TRYP-TOPHAN AMINOTRANSFERASE-RELATED (TAA1/TAR) family of aminotransferases (Stephanova et al., 2008). IPyA is subsequently converted into IAA by the enzymatic activity of the YUCCA (YUC) flavin-containing monooxygenases (Zhao et al., 2001). Expression of the bacterial tryptophan-2-monooxygenase (iaaM) auxin biosynthesis gene of under the control of a *YUC* promoter in *yuc* knock-out mutants (Cheng et al., 2006) or application of IAA (Chen et al., 2014) demonstrated the essential roles of the *YUC* genes in auxin biosynthesis in Arabidopsis. The IPyA pathway route is proposed as the main auxin biosynthesis route in Arabidopsis and has also been found in several other plant species, suggesting that it is a highly conserved IAA biosynthesis pathway in plants (Zhao, 2018).

Auxin is not produced in all plant cells, but is transported from source to sink tissues via the phloem or by polar cell-to-cell transport (Adamowski and Friml, 2015). This cell-to-cell transport of auxin is mainly mediated by plasma membrane-localized auxin efflux and influx carrier proteins. The PIN-FORMED (PIN) proteins have been identified as auxin efflux carriers that due to their asymmetric localization at the plasma membrane mediate unidirectional export of auxin from cells, thereby driving polar auxin transport (PAT) in plant tissues (Friml, 2010; Habets and Offringa, 2014). IAA can enter the cell by passive diffusion or through active import by influx carriers. The generally symmetrically localized AUXIN1/LIKE-AUX1 (AUX1/LAX) membrane proteins have been shown to act as permeases that mediate efficient auxin import (Péret et al., 2012; Swarup and Bhosale, 2019). AUX1/LAX-mediated IAA influx was shown to be 15 times more efficient than passive IAA diffusion and is therefore considered the predominant mode of auxin import in the cell (Swarup et al., 2005). The spatial localization pattern of PIN efflux carriers together with AUX1/LAX influx carriers determines the direction of auxin flow and differential accumulation of auxin in organs (Swarup and Bhosale, 2019).

The first phase of the plant life cycle starts with the fusion of the male and female gametes during fertilization to generate the zygote. This developmental switch, which is defined as gametophyte-to-zygotic transition, coincides with one of the most complex cellular reprogramming events, transforming the highly specialized, meiotically programmed egg cell into a totipotent mitotically active embryonic cell (She and Baroux, 2014). How the zygotic cell acquires totipotency remains largely unknown. During zygotic embryogenesis (ZE) the zygote gives rise to an embryo through cell division and morphogenesis. *De novo* auxin biosynthesis, auxin transport, and auxin signaling play critical roles in patterning and morphogenesis during ZE (Lau et al., 2012; Möller et al., 2017). The IPyA pathway regulates ZE by modulating spatiotemporal auxin production within embryonic tissues (Cheng et al., 2006; Stepanova et al., 2008). In Arabidopsis, the *YUC1*, *Y*UC*4*, *YUC8*, *YUC*9, *Y*UC10*, YUC11* genes are expressed in the 8- and 16-cell and globular embryo stages, and *yuc3 yuc9* and *yuc4 yuc9* double and *yuc1 yuc4 yuc10 yuc11* quadruple mutants exhibit embryonic patterning defects (Cheng et al., 2006; Robert et al., 2013). Recently, it has been shown that maternally biosynthesized auxin in the fertilized ovules also provides a source of auxin for the early-stage ZE (Robert et al., 2018).

Asymmetric distribution of auxin by PIN carriers also has been shown to play a vital role in Arabidopsis embryo patterning: i.e. the apical-basal and radial embryo axis formation and the establishment of bilateral symmetry by cotyledon initiation (Friml et al., 2003; Weijers et al., 2005). In addition to the PIN carriers, two auxin influx carriers LAX1 and AUX1 also contribute to formation the auxin gradients and auxin flow direction during ZE (Ugartechea-Chirino et al., 2010; Robert et al., 2015). The stronger embryo defects observed after combining mutations in *PIN* and *AUX1*/*LAX* genes indicate a cooperative function between auxin efflux and efflux carriers in controlling embryo development (Robert et al., 2015).

The ability of a plant cell to acquire totipotency and enter the embryogenesis program is not restricted to the zygote, as embryos can also develop from somatic ovule cells or unreduced gametophytes without fertilization in apomictic plant species (Ozias-Akins, 2006; Hand and Koltunow, 2014). In many flowering plants, vegetative somatic cells can also be converted to embryonic cells under appropriate *in vitro* conditions, in a process called somatic embryogenesis (SE). Besides providing a powerful tool for applications in plant biotechnology and plant breeding, including genetic transformation, somatic hybridization, clonal propagation and synthetic seed production, SE offers the potential for understanding cellular and molecular mechanisms that occur during plant embryo initiation and subsequent morphogenesis (Leljak-Levanić et al., 2015; Guan et al., 2016). Given the importance of SE for plant breeding and propagation, many attempts have been made to understand the molecular basis of this phenomenon in different plant species. Some genes have been identified that encode transcription factors that promote SE. Ectopic expression of a single gene like *BABY BOOM* (*BBM*), *LEAFY COTYLEDON 1* (*LEC1*), *LEC2*, *WUSCHEL* (*WUS*) or *AT-HOOK MOTIF NUCLEAR LOCALIZED 15* (*AHL15*) induces spontaneous SE (Lotan et al., 1998; Stone et al., 2001; Boutilier et al., 2002; Zuo et al., 2002; Karami et al., 2021). SE can also be achieved by exogenous application of plant hormones. Recent research has provided new insights into the process of transcription factor- or hormone-induced SE (Horstman et al., 2017; Wójcik et al., 2020), but still the developmental, hormonal and molecular mechanisms governing SE are complex and far from understood.

Sixty-five percent of the recent SE protocols use the herbicide 2,4-dichlorophenoxyacetic acid (2,4-D), a synthetic analog of the natural auxin IAA, for SE induction (Wójcik et al., 2020). Although 2,4-D mimics IAA at the molecular level, 2,4-D is much more stable in plant cells than IAA (Eyer et al., 2016). Several studies have shown that 2,4-D or other exogenously-applied auxins significantly increase the level of IAA in the explants undergoing SE (Ivanova et al., 1994; Michalczuk and Druart, 1999; Jiménez and Bangerth, 2001a; Jiménez and Bangerth, 2001b; Cheng et al., 2016; Márquez-López et al., 2018; Vondrakova et al., 2018; Awada et al., 2019). IAA accumulation was also found in embryogenic tissues induced by *LEC2* overexpression in Arabidopsis seedlings (Stone et al., 2008). In Arabidopsis, 2,4-D induces expression of several *YUC* genes in IZE explants and embryogenic callus, and higher order *yuc* mutants produce fewer somatic embryos per explant compared to wild-type explants (Bai et al., 2013; Wójcikowska et al., 2013). In addition, ethylene has been reported to have a negative impact on SE in Arabidopsis by reducing *YUC* expression and thereby lowering the auxin levels in embryogenic callus (Bai et al., 2013). Therefore, IAA biosynthesis in embryonic cells seems to plays a significant role in somatic embryo induction. However, the exact action of endogenous auxin in the early stage of somatic embryo induction has not been well characterized.

Here we use live-cell imaging and chemical biology approaches to reveal new insights into the contribution of endogenous auxin action in both 2,4-D and *AHL15* gene induced SE. Our data show that induction of embryonic stem cells from somatic explants does not require endogenous auxin biosynthesis, whereas an increase in endogenous auxin levels in conjunction with auxin transport is essential to maintain embryo identity and promote embryo differentiation.

## Results

### The *pWOX2:NLS-YFP* reporter distinguishes different stages of SE in Arabidopsis

Immature zygotic embryos (IZEs) from Arabidopsis are a much used experimental system to study SE induction in response to 2,4-D (Gaj, 2001). In our hands, embryonic callus can be efficiently induced on cotyledons of IZEs incubated for seven to nine days of culture on medium supplemented with 4.5 μM 2,4-D. Following transfer of the explants to 2,4-D free medium, this embryonic callus develops into globular and further into cotyledon-stage somatic embryos (Ikeda-Iwai, 2002; Gaj, 2011). Recently, we showed that overexpression of the *AHL15* gene also induces SE on cotyledon tissue of IZEs in the absence of 2,4-D (Karami et al., 2021). In *p35S:AHL15* cotyledons, the protodermal cells at the adaxial side are converted into embryonic callus around six days after culture. Approximately two days later these embryonic cells develop into globular shaped pro-embryos.

Here we used the *pWOX2:NLS-YFP* reporter (Breuninger et al., 2008) for time-lapse imaging of embryo initiation during 2,4-D- and *AHL15*-induced SE. *WOX2* is a member of the *WUSCHEL* (*WUS*) homeodomain gene family and the reporter is expressed in the Arabidopsis zygote, the suspensor and the early zygotic embryo (Supplemental Figure 1A). In our 2,4-D-based SE system, expression of the *pWOX2:NLS-YFP* reporter was not detectable in IZE cotyledons within the first five days of IZE culture (Supplemental Figure 1B). Relatively weak *pWOX2:NLS-YFP* activity was first detected in the adaxial regions of cotyledons after six to seven days. One to two days later, this signal increased in the areas that formed embryogenic protrusions (Supplemental Figure 1B). No *pWOX2:NLS-YFP* activity was detected in wild-type IZE cotyledons cultured in the absence of 2,4-D (Supplemental Figure 1B). Also in *p35S:AHL15* IZEs, the *pWOX2:NLS-YFP* signal was not detected in the cotyledons of *p35S:AHL15* explants within the first four days of culture (Supplemental Figure 2A). After 5 to 6 days, a relatively weak *pWOX2:NLS-YFP* signal was observed in epidermal cells at the adaxial side of *p35S:AHL15* IZE cotyledons, just like with the 2,4-D system (Supplemental Figure 2B). One to two days later, *pWOX2:NLS-YFP* expression significantly increased (Supplemental Figure 2C), followed by a reduction in expression in developing globular embryos on day nine to 11 (Supplemental Figure 2D,E). Thus *pWOX2:NLS-YFP* is not expressed in cotyledon somatic cells within the first four days of culture, but is induced later, becomes highly expressed in dividing embryonic clusters and is then down-regulated from the globular stage onward (Supplemental Figure 1C). The results indicate that *WOX2* is a good marker for both our 2,4-D- and AHL15-induced SE systems to identify cell fate transitions to embryo development and for marking the developmental stages of SE.

Based on these observations, we defined three distinct developmental stages during the early process of SE induction from Arabidopsis IZEs: 1) acquisition of embryogenic competence in somatic cells around day 6, 2) rapid cell proliferation coinciding with the conversion of competent somatic cells into embryonic stem cells around day 8 and 3) the development of embryonic cells into globular pro-embryos around day 10 of culture (Supplemental Figure 1C).

### The IPyA auxin biosynthesis pathway is essential for 2,4-D-induced SE

The YUC flavin monooxygenases catalyze the rate-limiting step in the main auxin biosynthesis pathway in plants (Zhao, 2018). Previous studies have shown that the *YUC1*, *Y*UC*2*, *YUC4*, *YUC*6, *YUC10, YUC11* genes are expressed at the sites of embryo formation during 2,4-D-induced somatic embryogenesis in Arabidopsis, suggesting that they are responsible for increased IAA biosynthesis in these embryogenic cells (Bai et al., 2013; Wickramasuriya and Dunwell, 2015). However, how an increase in IAA biosynthesis affects the progression of 2,4-D-induced SE is not clear. To determine the role of endogenous IAA biosynthesis in 2,4-D-induced SE, we first re-analyzed the activity of several *pYUC*:*GFP-GUS* reporters in IZE explants on days 0, 3, 5 and 7 of culture in medium supplemented with 2,4-D (Supplemental Figure 3). The *pYUC2/10:GFP-GUS* reporters were not or only barely expressed at any time point in culture (not shown), but dynamic expression patterns were observed for the *pYUC4/5/6/7/8/9/11:GFP-GUS* reporters (Supplemental Figure 3). The reporters for *YUC6/7/8/9* were strongly expressed in cotyledon tissue in seven-day-old IZE explants cultured on 2,4-D medium, whereas they barely showed expression in seedlings-derived from IZE explants that were cultured on medium lacking 2,4-D (Figure 1A). These *YUC* genes might be responsible for the increase in the auxin biosynthesis in embryogenic tissues induced on cotyledons.

**Figure 1.**
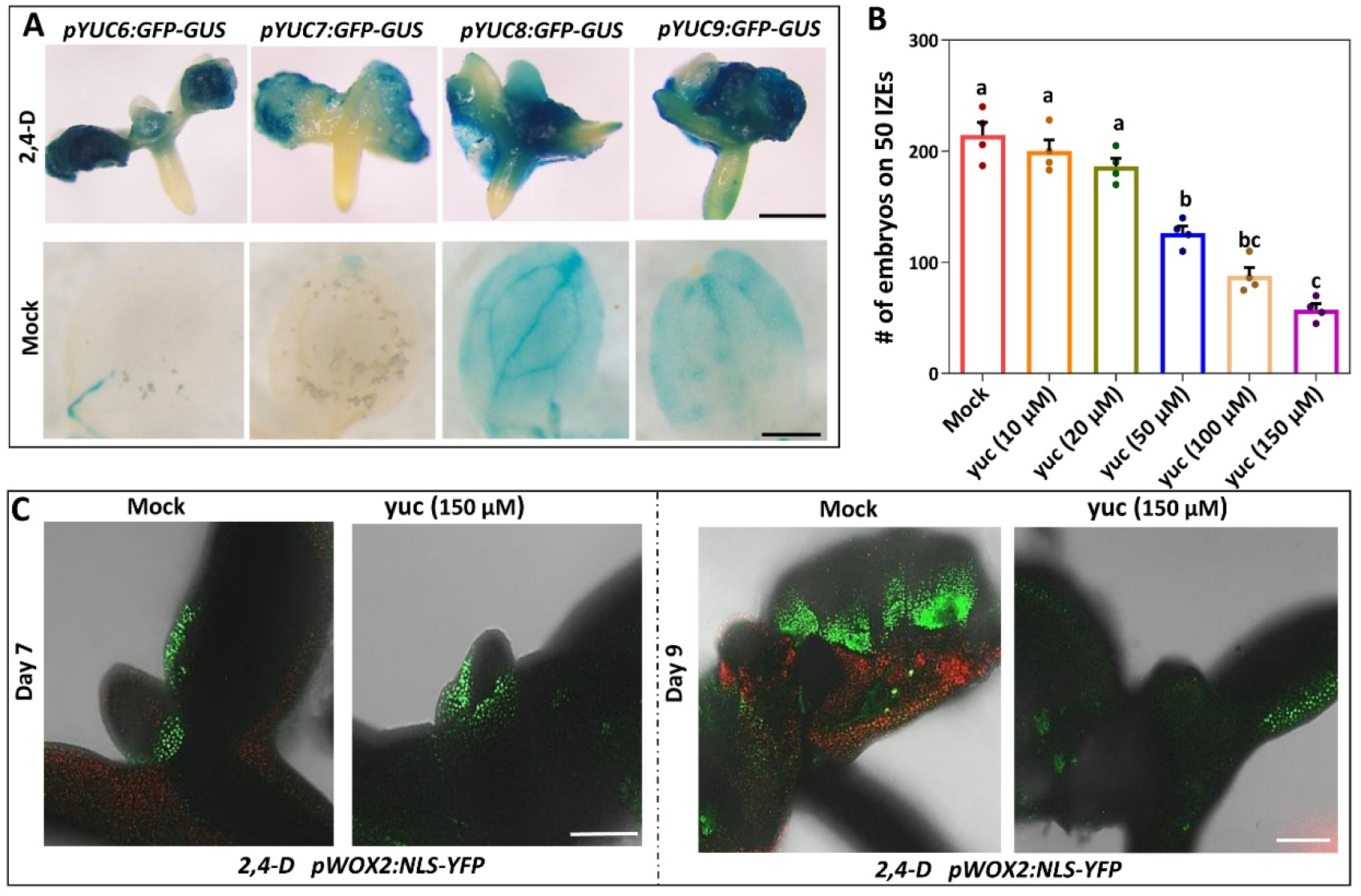
IAA biosynthesis in cotyledon tissues is essential for 2,4-D induced-SE. (**A**) Expression pattern of *pYUC6:GFP-GUS*, *pYUC7:GFP-GUS*, *pYUC8:GFP-GUS*, and *pYUC9:GFP-GUS* reporters in immature zygotic embryos (IZEs) cultured for 8 days on medium with 2,4-D (upper panel) or in cotyledons of IZEs germinated for 8 days on medium without 2,4-D (lower panel). (**B**) Effect of different concentrations of the auxin biosynthesis inhibitor yucacin (yuc) on the capacity to induce somatic embryos on IZEs cultured on medium with 2,4-D. Dots indicate in the number somatic embryos produced on cotyledons of 50 IZEs (n=4 biological replicates), bars indicate the mean and error bars the s.e.m.. Different letters indicate statistically significant differences (P < 0.001) as determined by a one-way analysis of variance with Tukey’s honest significant difference post hoc test. (**C**) Expression of *pWOX2:NLS-YFP* in cotyledons of wild-type IZEs cultured for seven (left) and nine (right) days on 2,4-D medium without (Mock) and with 150 μM yuc. Size bars indicate 1 mm.

To confirm the role of IPyA pathway-mediated auxin biosynthesis in 2,4-D-induced SE and in view of the strong redundancy between *YUC* genes during ZE and SE(Cheng et al., 2007; Bai et al., 2013; Robert et al., 2013). we used yucasin (yuc), a specific inhibitor of YUC enzyme activity (Nishimura et al., 2014), to reduce IAA levels during embryogenic callus induction by 2,4-D. Treatment with 50 μM or higher concentrations of yuc resulted in a significant reduction in the number of somatic embryos (Figure 1B). We propose that the lack of a significant effect of lower yuc concentrations (10 and 20 μM, Figure 1B) is due to the higher auxin level that is already present in 2,4-D-induced explants.

Unexpectedly, we observed *pWOX2:NLS-YFP* expression in both untreated and yuc-treated cotyledons starting from six to seven-day-old 2,4-D-treated IZE explants (Figure 1C). However, whereas *pWOX2:NLS-YFP* expression increased in untreated explants one to two days later, it decreased in cotyledons of yuc-treated explants (Figure 1C). These results indicate that endogenous auxin is not required for the initiation of SE, but rather for the maintenance of embryonic cell identity in 2,4-D-induced SE.

### Auxin biosynthesis by the IPyA pathway is also essential for *AHL15*-induced SE

Next we studied the role of auxin biosynthesis in *AHL15-*induced SE. As gene-induced SE occurs in the absence of exogenous 2,4-D, this allowed us to follow the spatiotemporal endogenous auxin dynamics in *p35S:AHL15* cotyledon tissues using the auxin responsive *pDR5:GFP* reporter (Benkova et al., 2003). Time-lapse analysis showed that *pDR5:GFP* activity was not different in wild-type and *p35S:AHL15* cotyledons during the first three days of culture (Figure 2A). One to two days later, however, reporter expression markedly increased throughout the entire *p35S:AHL15* cotyledons, while no or a much lower GFP signal was observed in wild-type explants (Figure 2A). These results suggested an increase in auxin levels in *p35S:AHL15* cotyledons.

**Figure 2.**
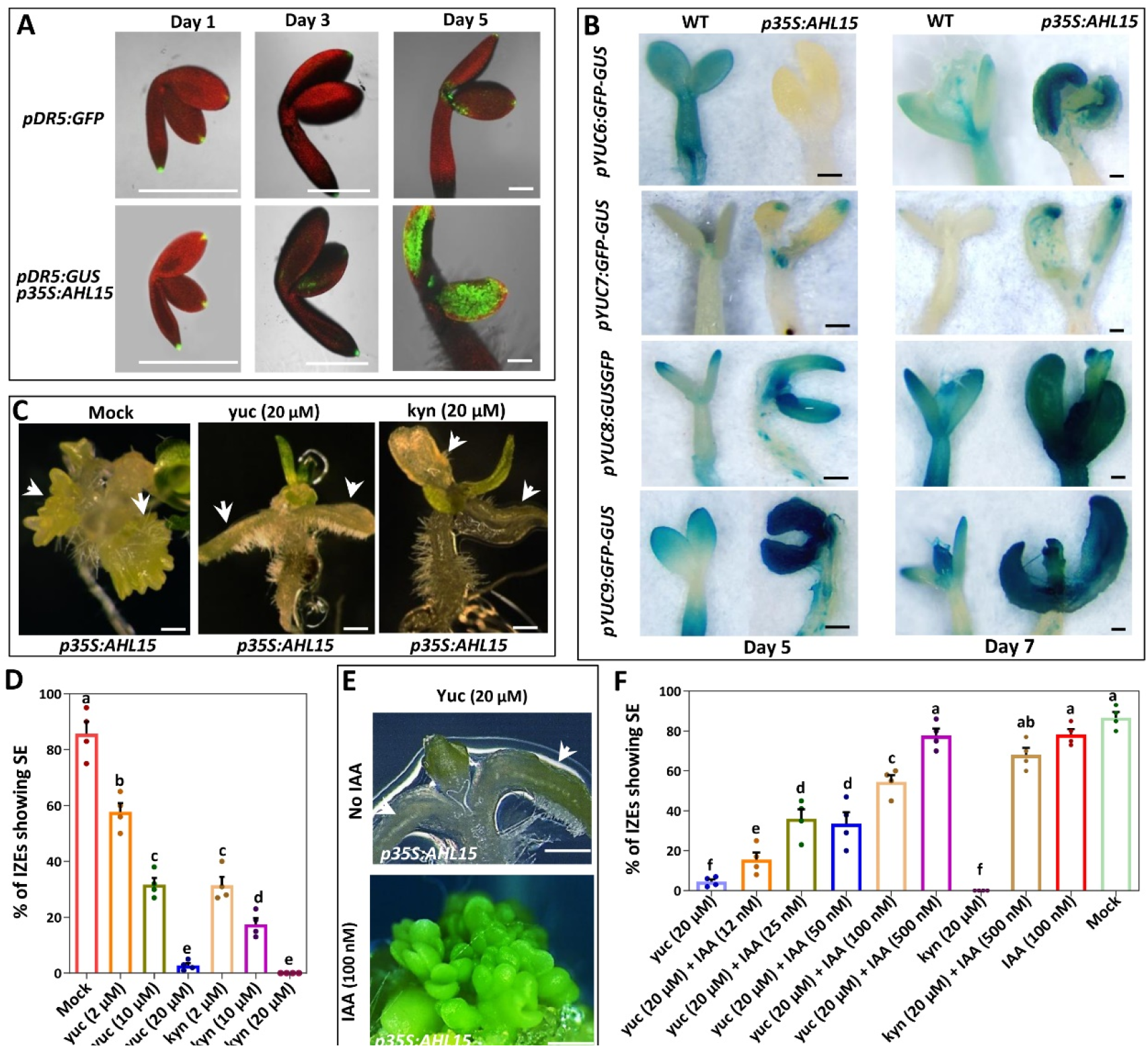
IAA biosynthesis in *35S:AHL15* cotyledon tissues is essential for AHL15-induced SE. (**A**) Expression of the *pDR5:GFP* reporter in wild-type (upper panel) or *p35S:AHL15* (lower panel) IZEs cultured for one, three or five days on medium without 2,4-D. (**B**) Expression pattern of *pYUC6:GFP-GUS*, *pYUC7:GFP-GUS*, *pYUC8:GFP-GUS* or *pYUC9:GFP-GUS* reporters in wild-type and *35S:AHL15* IZEs cultured for five (right) or seven (left) days on medium without 2,4-D. (**C**) The phenotypes of *p35S:AHL15* IZEs cultured for 2 weeks on medium without 2,4-D and without (Mock) or with 20 μM of the auxin biosynthesis inhibitor yuc (middle) or kyn (left). White arrowheads indicate the adaxial side of cotyledons. (**D**) Effect of different concentrations of yuc and kyn on the efficiency of somatic embryo induction on cotyledons of *p35S:AHL15* IZEs. (**E**) The phenotypes of *p35S:AHL15* IZEs cultured for two weeks on medium without 2,4-D, but with 20 μM yuc and without (No IAA) or with 100 nM IAA. White arrowheads indicate adaxial side of cotyledons. (**F**) Exogenous IAA treatment restores yuc- or kyn-impaired SE on cotyledons of *p35S:AHL15* IZEs. Dots in **D** and **F** indicate the percentage of *p35S:AHL15* IZEs producing somatic embryos (n=4 biological replicates, with 50 IZEs per replicate), bars indicate the mean, error bars the s.e.m. and different letters indicate statistically significant differences (P < 0.001) as determined by a one-way analysis of variance with Tukey’s honest significant difference post hoc test. Size bars in **A**, **B**, **C** and **E** indicate 1 mm.

Comparison of the expression of *pYUC:GFP-GUS* and *pYUC:NLS-3xGFP* reporters in *p35S:AHL15* or wild-type IZE cultures at five and seven days of cultures did not detect obvious differences in activity of the *YUC1/2/3/4/5/10/11* promoters (not shown), indicating that the increase in *pDR5* expression in *p35S:AHL15* cotyledons is not likely to be mediated by these *YUC* genes. At the same time points the *pYUC7/8/9* promoters did show higher activity in *p35S:AHL15* compared to wild-type explant cotyledons (Figure 2B). Interestingly, in five-day-old IZE explants, *pYUC6:GFP-GUS* expression was not detected in *35S:AHL15* cotyledons, whereas it was expressed in wild-type explants (Figure 2B). After 7 days of culture, however, expression of this reporter was strongly upregulated in *p35S:AHL15* cotyledons (Figure 2B), whereas it was reduced in wild-type cotyledons. These results suggested that the increase in *pDR5* expression in *p35S:AHL15* cotyledons is caused by the induction of *YUC6/7/8/9* gene expression.

We did not detect a significantly higher level of *pDR5:GFP* activity or *pYUC6/7/8/9* reporter expression in hypocotyl or root tissues of *p35S:AHL15* compared to wild-type explants (not shown). This implies that *YUC6/7/8/9* genes are specifically upregulated in cotyledon tissues, causing a cotyledon-specific increase in auxin response in *p35S:AHL15* explants. The simultaneous induction of *pWOX2:NLS-GFP* reporter expression in *p35S:AHL15* explant cotyledons (Figure 3A) suggested that the enhanced auxin biosynthesis and response is associated with either the cell fate change or the acquisition of embryo identity in cotyledons tissues.

**Figure 3.**
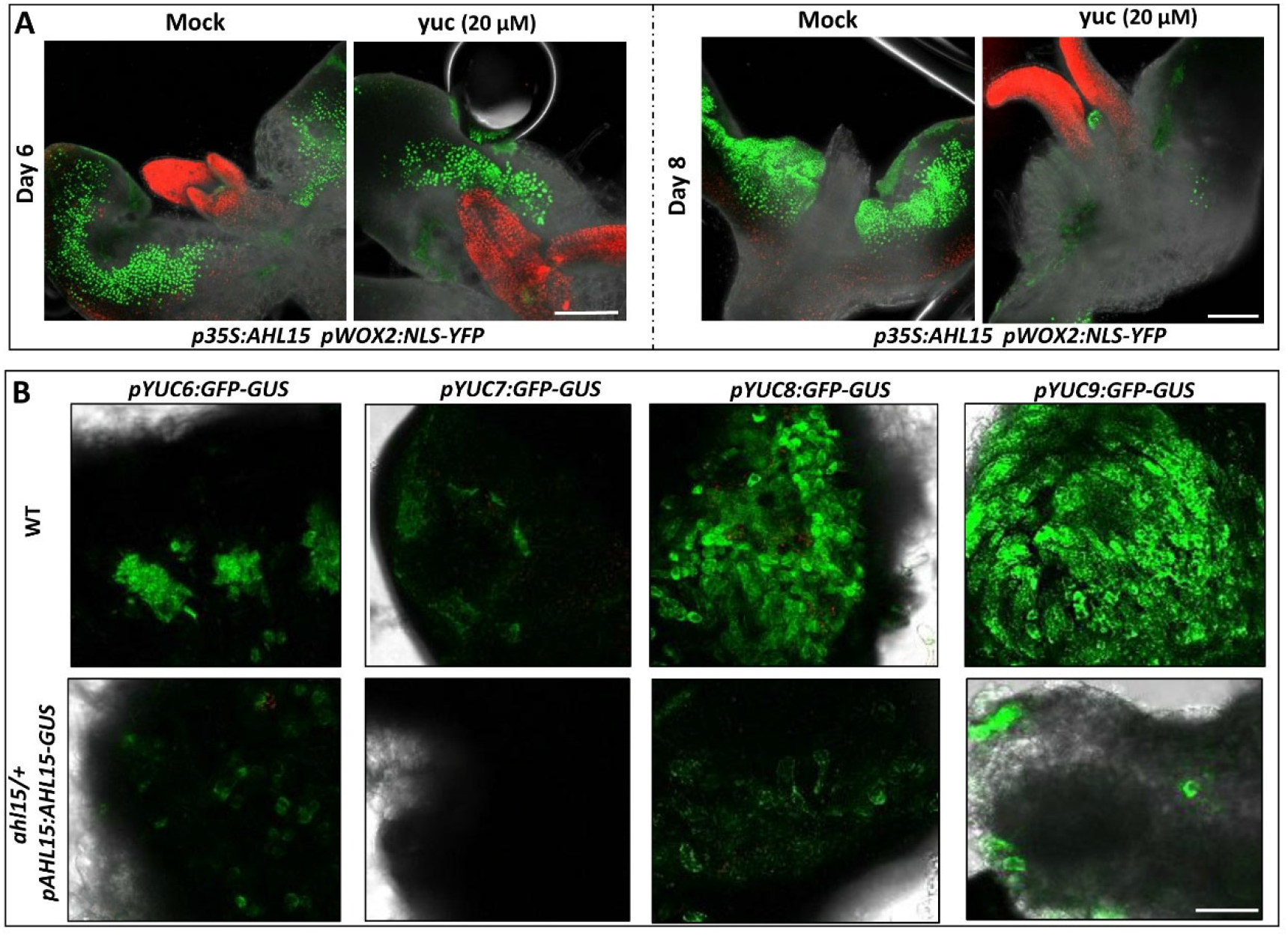
YUC-mediated auxin biosynthesis is required for the maintenance of embryonic cell identity. (**A**) The expression of *pWOX2:NLS-YFP* in cotyledons of germinating *p35S:AHL15* IZEs after six (left) and eight (right) days of culture on medium without (Mock) or with 20 μM yuc. (**B**) Expression of the *pYUC6:GFP-GUS*, *pYUC7:GFP-GUS*, *pYUC8:GFP-GUS* or *pYUC9:GFP-GUS* reporters in cotyledons of wild-type (WT) or *ahl15/+ pAHL15:AHL15-GUS* IZEs cultured for eight days on 2,4-D medium. Size bars indicate 0.5 mm.

Treatment with the yuc auxin biosynthesis inhibitor or with L-kynurenine (kyn), which specifically blocks the tryptophan aminotransferase TAA1/TAR enzymes of the IPyA pathway (He et al., 2011), completely inhibited somatic embryo induction in *p35S:AHL15* IZE explants. Inhibition occurred at relatively low (20 μM) yuc or kyn concentrations (Figure 2C and D), compared to 2,4-D-induced SE, and could be rescued by exogenous application of 12 to 500 nM IAA, concentrations that did not inhibit SE in the absence of yuc or kyn (Figure 2E and F). These results, together with the observed induction of *YUC6/7/8/9* gene expression, suggest that the IPyA pathway-mediated IAA biosynthesis is required for *AHL15*-induced SE. The rescue of yuc- or kyn-inhibited SE by exogenous application of relatively low auxin concentrations (Figure 2F) suggests that AHL15-induced SE is hypersensitive to changes in auxin levels, implying that a relatively small increase in auxin levels is sufficient for AHL15-induced SE.

Expression of the *pWOX2:NLS-YFP* reporter was detected in the cotyledons of both five to six-day-old untreated and yuc-treated *p35S::AHL15* IZE explants (Figure 3A). However, one to two days later, when *pWOX2:NLS-YFP* expression increased in the untreated control, *pWOX2:NLS-YFP* expression decreased or disappeared in cotyledons of yuc-treated *p35S::AHL15* IZEs (Figure 3A). These results indicate that the cotyledon cells in yuc-treated *p35S:AHL15* IZE explants initially acquire embryo identity, but that these freshly induced-embryonic cells are not stable and quickly return to the non-embryogenic state in the absence of auxin biosynthesis. Thiswas also observed for 2,4-D-induced SE and implies that endogenous auxin production mediated by the IPyA pathway is not required for the acquisition of embryo identity, but that it mainly contributes to the maintenance of embryonic identity and for embryo development.

We previously reported that *AHL15* is highly expressed in 2,4-D-induced embryogenic tissues and that expression of a *pAHL15:AHL15-GUS* fusion in the *ahl15*/+ heterozygous or *ahl15* homozygous mutant background respectively inhibits 2,4-D-induced SE or arrests zygotic embryogenesis. Expression of the AHL15-GUS fusion in these mutant backgrounds leads to a dominant-negative effect that overcomes the functional redundancy between *AHL15* and other *AHL* family members, and thus leads to *ahl* loss-of-function (Karami et al., 2021). The expression of *YUC6/7/8/9* was significantly reduced in cotyledons of 2,4-D-treated *ahl15/+ pAHL15:AHL15-GUS* IZEs compared to wild-type IZEs (Figure 3B), suggesting that these *YUC* genes may act downstream of AHL15 during 2,4-D-induced SE. The significantly lower expression of the *pDR5:GFP* reporter in *ahl15 pAHL15:AHL15-GUS* zygotic embryos (Supplemental Figure 4B) compared to wild-type embryos (Supplemental Figure 4A) suggests that *AHL* genes also play a role in stimulating IAA biosynthesis in zygotic embryos, possibly through induction of *YUC* gene expression.

### Auxin efflux is not required for SE initiation but for proper embryo patterning

Auxin efflux carriers play an important role in zygotic embryo patterning, but not in the initiation of ZE (Friml et al., 2003). In our hands, 2,4-D-induced SE is a clear two-step process, involving 1) induction of embryonic callus on the IZE explant cotyledons after about ten days of culture on 2,4-D containing medium, and 2) patterning of this embryonic callus into somatic embryos after explant transfer to 2,4-D free medium. Obviously, the patterning process is inhibited by the presence of exogenous 2,4-D. To determine at which stage of SE auxin efflux is important, we analyzed the effect of the auxin efflux inhibitor N-1-naphthylphthalamic acid (NPA) on these two steps of 2,4-D-induced SE. Our experiments showed that the number of somatic embryos is only slightly decreased following treatment with different concentrations of NPA during the first step (initiation) of 2,4-D-induced SE (Figure 4A and B). By contrast, NPA treatment led to aberrant embryo-like structures (Figure 4A) and strongly reduced the number of normal, non-fused somatic embryos produced when applied during the second (patterning) step of 2,4-D-induced SE (Figure 4C). As we cannot exclude that the slight effect of NPA treatment during the first step is caused by NPA accumulation persisting during the second step, we conclude that auxin efflux plays no or only a minor role in the initiation of SE, but that like in ZE it has a major role later in embryo patterning and development.

**Figure 4.**
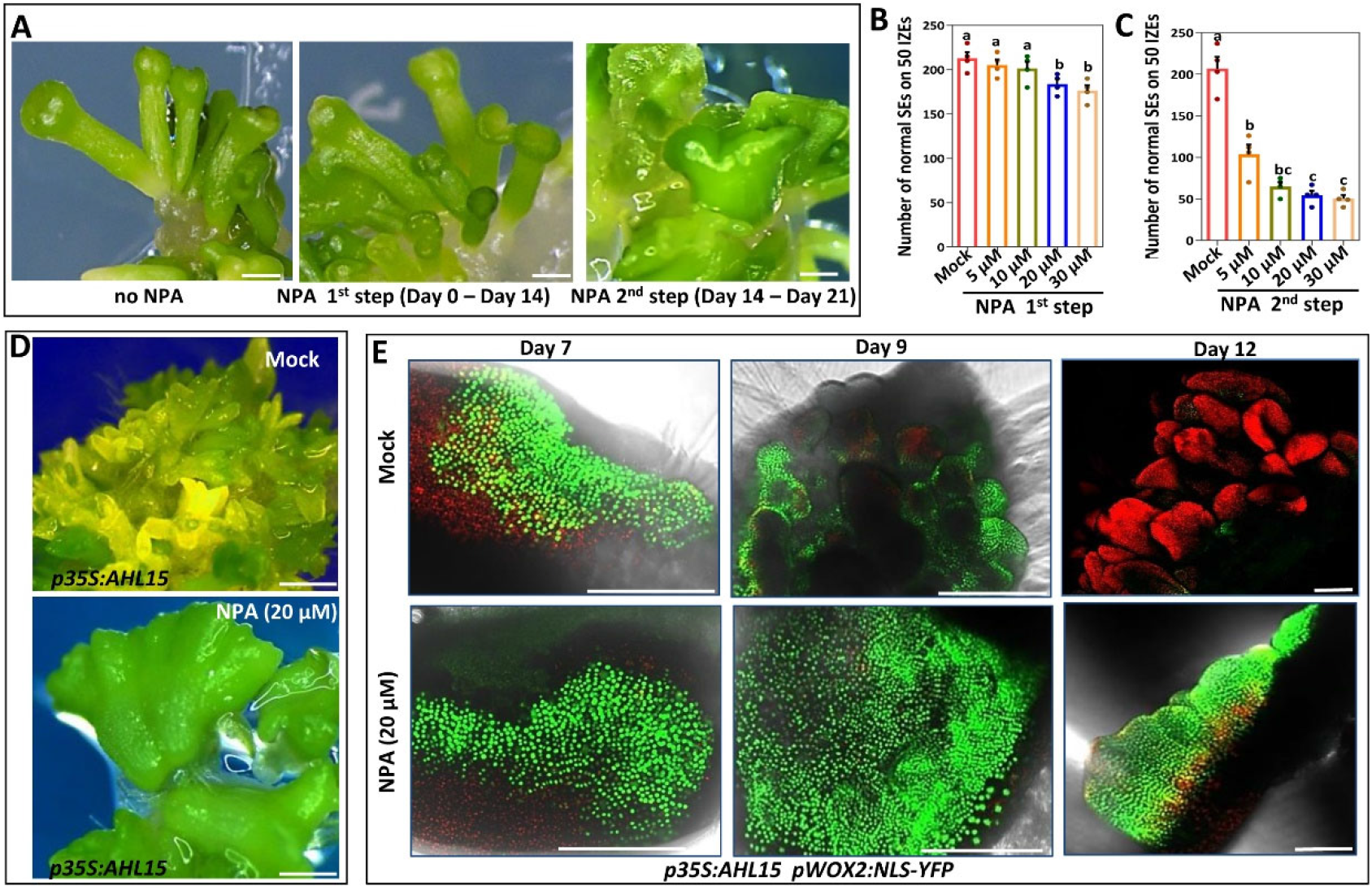
Auxin efflux is required for the proper development of embryonic cells into somatic embryos. (**A**) The phenotype of somatic embryos formed on cotyledons of wild-type IZEs that were first grown for two weeks on 2,4-D medium and subsequently cultured for 1 week on medium without 2,4-D (left), or with 20 μM NPA (middle) or first on 2,4-D medium with 20 μM NPA and subsequently cultured on medium without 2,4-D and NPA (right). (**B**) The number of non-fused somatic embryos (normal SEs) per 50 IZEs that were first grown for two weeks on 2,4-D medium without (Mock) and with different concentrations of NPA, and subsequently grown for 1 week on medium without 2,4-D or NPA. (**C**) The number of non-fused somatic embryos (normal SEs) per 50 IZEs that were first grown for 2 weeks on 2,4-D medium and subsequently grown for 1 week on medium without 2,4-D and with different concentrations of NPA. The dots in **B** and **C** indicate the number of normal somatic embryos produced per 50 IZEs (n=4 biological replicates), bars indicate the mean and error bars indicate s.e.m.. Different letters indicate statistically significant differences (P < 0.001) as determined by one-way analysis of variance with Tukey’s honest significant difference post hoc test. (**D**) The phenotypes of somatic embryos formed on cotyledons of a two week-old *p35S:AHL15* IZE on B5 medium supplemented with 20 μM NPA (right) and without NPA (left). (**E**) The expression pattern of *pWOX2:NLS-YFP* in cotyledon tissues of *p35S:AHL15* IZEs after seven, nine, or twelve days of culture on medium without NPA (upper images) or on medium supplemented with 20 μM NPA (lower images). Size bars indicate 1 mm in **A**, **B**, **D**, and **E**, or 0.5 mm in **E**.

As *AHL15*-induced SE occurs in the absence of exogenous 2,4-D, the patterning process is not inhibited, and embryo initiation and patterning occur more simultaneous compared to 2,4-D-induced SE. Therefore, the effect of NPA on *AHL15*-induced SE was only tested from the start of *p35S:AHL15* IZE culture. NPA-treated *p35S:AHL15* IZEs only developed a few aberrant embryos with fused cotyledons (Figure 4D), whereas many somatic embryos (around ten to twenty per explant) were formed without NPA treatment (Figure 4D). This result reveals that auxin efflux is required for the efficient production and the proper development of somatic embryos during *AHL15*-induced SE.

Expression of the *pWOX2:NLS-YFP* embryo identity reporter in *p35S:AHL15* IZE cotyledons after 7 days of culture was similar in the presence or absence of NPA (Figure 4E). Without NPA, we observed the usual reduction in *pWOX2:NLS-YFP* expression during further patterning and development of somatic embryos after 9 and 12 days of culture (Figure 4E). In the presence of NPA, however, we did not observe this reduction in *pWOX2:NLS-YFP* expression (Figure 4E). High expression of *pWOX2:NLS-YFP* persisted until day 12 in NPA cultured *AHL15-*induced somatic embryos, which might reflect the maintenance of early embryo identity and inhibition of subsequent embryo patterning and development, probably caused by intracellular auxin accumulation due to lack of efflux, similar to what is observed for the 2,4-D-cultured embryonic calli. Taken together these data indicate that in both 2,4-D- and *AHL15*-induced SE, the early events, i.e. acquisition and maintenance of embryonic identity and the induction of embryonic cells, require high intracellular auxin and can proceed independently of auxin efflux, while the subsequent embryo patterning and development relies on auxin efflux to reduce the intracellular auxin concentration allowing loss of embryo identity. Polarly localized PIN proteins on the plasma membrane are known to be key components driving auxin efflux-mediated patterning during zygotic embryogenesis (Friml et al., 2003). Therefore, we examined the expression of *pPIN1:PIN1-GFP*, *pPIN2:PIN2-VENUS, pPIN4:PIN4-GFP*, and *pPIN7:PIN7-GFP* reporters during SE. Of these reporters, only PIN1-GFP expression was observed in *35S:AHL15* (Figure 5A) or 2,4-D-cultured IZE cotyledons (Figure 5B). The earliest PIN1-GFP signals were detected after seven to eight days of culture at the abaxial side of the cotyledons (Figure 5A and B). These results indicate that PIN1 is the major regulator of auxin efflux during AHL15 and 2,4-D-induced SE.

**Figure 5.**
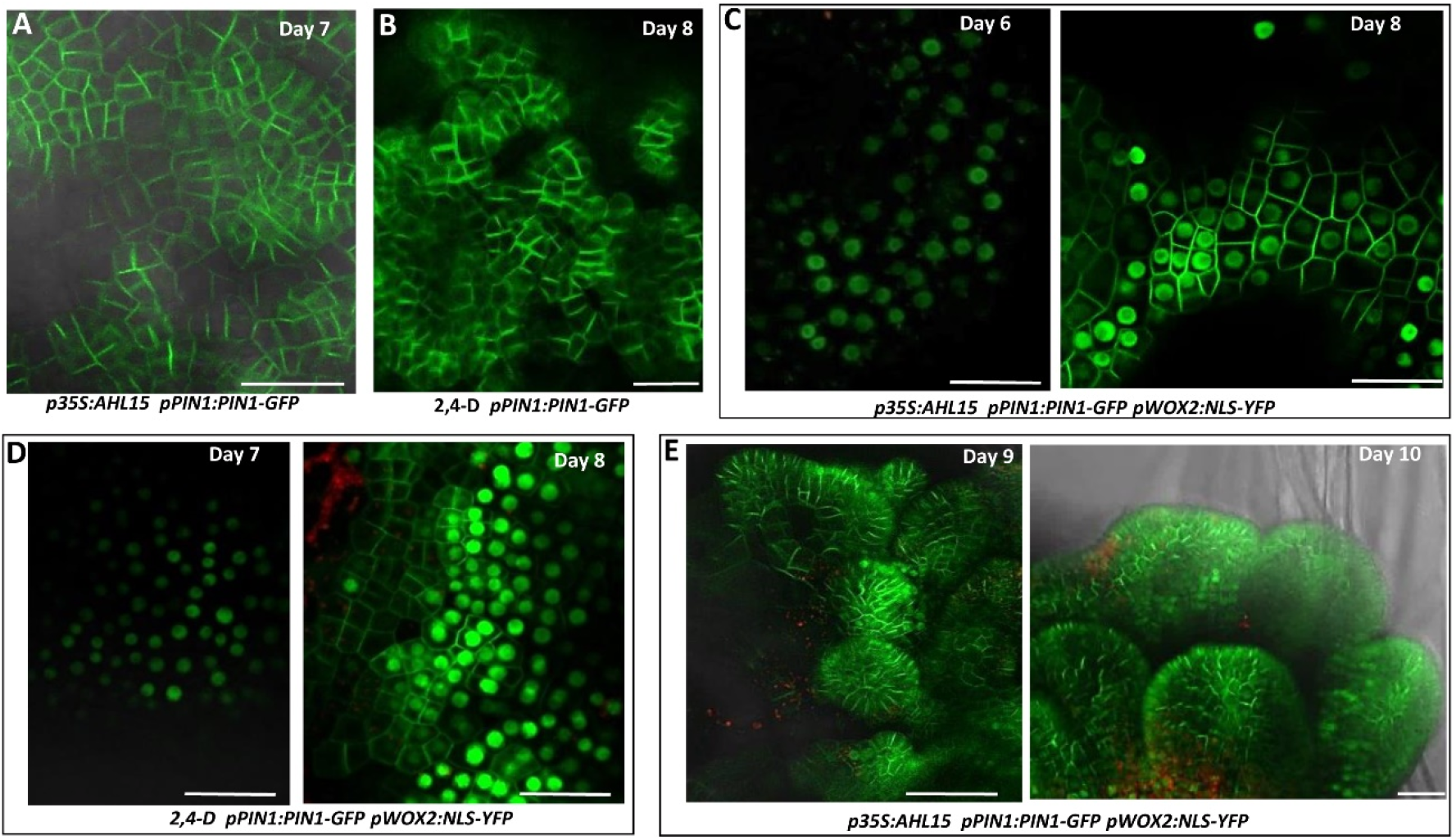
Expression and localization of *PIN1* during AHL15- and 2,4-D-induced SE. (**A, B**) PIN1-GFP signals detected in cotyledons of *p35S:AHL15* IZEs after seven days of culture on medium (A) or in cotyledons of wild-type IZEs after eight days of culture on medium supplemented with 5 μM 2,4-D (B). (**C**) Expression of *pIN1:PIN1-GFP* (plasma membrane) and *pWOX2:NLS-YFP* (nucleus) in cotyledon tissues of *p35S:AHL15* IZEs after six (left) or eight (right) days of culture. (**D**) Expression of *pPIN1:PIN1-GFP* and *pWOX2:NLS-YFP* in cotyledon tissues of wild-type IZEs after seven (left) or eight (right) days of culture on medium supplemented with 5 μM 2,4-D. Note that in **C** (left) and **D** (left) the cotyledon cells show a clear nuclear YFP signal, whereas no PIN1-GFP is yet detectable. (**E**) Expression of *pPIN1:PIN1-GFP* and *pWOX2:NLS-YFP* in globular (left) and heart (right) stage embryos developing on cotyledons of *p35S:AHL15* IZEs cultured for respectively nine or ten days. Size bars indicate 100 μm.

To further monitor PIN1 activity during SE, co-expression of *pWOX2:NLS-YFP* and *pPIN1:PIN1-GFP* were tracked in the *AHL15-* and 2,4-D-induced SE systems. Time-lapse experiments showed that *pPIN1:PIN1-GFP* is not expressed when the first *pWOX2:NLS-YFP* activity appears in the AHL15*-* and 2,4-D-induced SE systems (Figure 5C and D). However, one to two days later, clear *pWOX2:NLS-YFP* and *pPIN1:PIN1-GFP* co-expression was detected in both systems (Figure 5C and D). During somatic embryo development, *pPIN1:PIN1-GFP* expression was maintained in the embryo, but *pWOX2:NLS-YFP* disappeared (Figure 5E). These results suggest that induction of embryonic cell identity is independent of PIN1 function, but that PIN1 promotes the development of embryonic cells toward multicellular embryos.

### Auxin influx is required for embryonic cell identity maintenance during SE

Auxin influx carriers facilitate the import of auxin into plant cells and thereby play a critical role in the directional auxin flow and the resulting auxin maxima and minima formed during ZE (Ugartechea-Chirino et al., 2010; Robert et al., 2015; Boot et al., 2016). When the auxin influx inhibitor 1-naphthoxyacetic acid (1-NOA) (Parry et al., 2001) was applied during the first step (initiation) of 2,4-D-induced SE, it strongly reduced the number of embryos formed on cotyledons (Figure 6A), suggesting that auxin influx is essential for this phase of SE. As with yuc treatment, AHL15-induced SE was significantly more sensitive to 1-NOA treatment. 30 μM 1-NOA completely blocked the induction of embryos on cotyledons of *p35S:AHL15* IZEs (Figure 6B), whereas this was not the case for 2,4-D-induced SE. To further explore whether impaired auxin influx affects the initiation or maintenance of embryonic cell identity during SE, *pWOX2:NLS-YFP* expression was tracked in 1-NOA-treated IZE explants. *pWOX2:NLS-YFP* expression was initially detected after six to seven days of culture in *35:AHL15* (Figure 6C) and 2,4-D treated IZE cotyledons (Figure 6D) in the presence or absence of 1-NOA. One to two days later, however, the *pWOX2:NLS-YFP* signals were highly reduced in cotyledons of 1-NOA-treated IZEs compared with mock-treated IZEs (Figure 6C and D). These results suggest that auxin influx is required for embryonic identity maintenance, just like auxin biosynthesis by the IPyA pathway. The negative effect of the yuc auxin biosynthesis inhibitor on *p35S:AHL15* IZEs could be complemented by providing exogenous IAA (Figure 2F). However, co-treatment of *p35S:AHL15* IZEs with 30 μM 1-NOA, 20 μM yuc, and 100 nM IAA disrupted this IAA-mediated complementation (Supplemental Figure 5), confirming that embryonic cell identity maintenance relies on elevated intracellular IAA levels mediated by both auxin biosynthesis and auxin influx. Analysis of *pAUX1:AUX1-GFP* and *pLAX1:LAX1-GFP* cultured IZEs showed that *LAX1* is not expressed during *AHL15* or 2,4-D induced SE (not shown), but that AUX1-GFP signals coincide with the appearance of *pWOX2:NLS-YFP* marked embryonic cells in cotyledons at day six to eight and are also present in globular embryos at day 10 (Figure 6E and F). These results suggest that AUX1 mediates auxin uptake during SE. The AUX1-GFP signals appear at the same time as the PIN1-GFP signals, suggesting that auxin influx and efflux balance the auxin levels in cells and thereby determine whether embryonic cell identity is maintained or that patterning and development are initiated.

**Figure 6.**
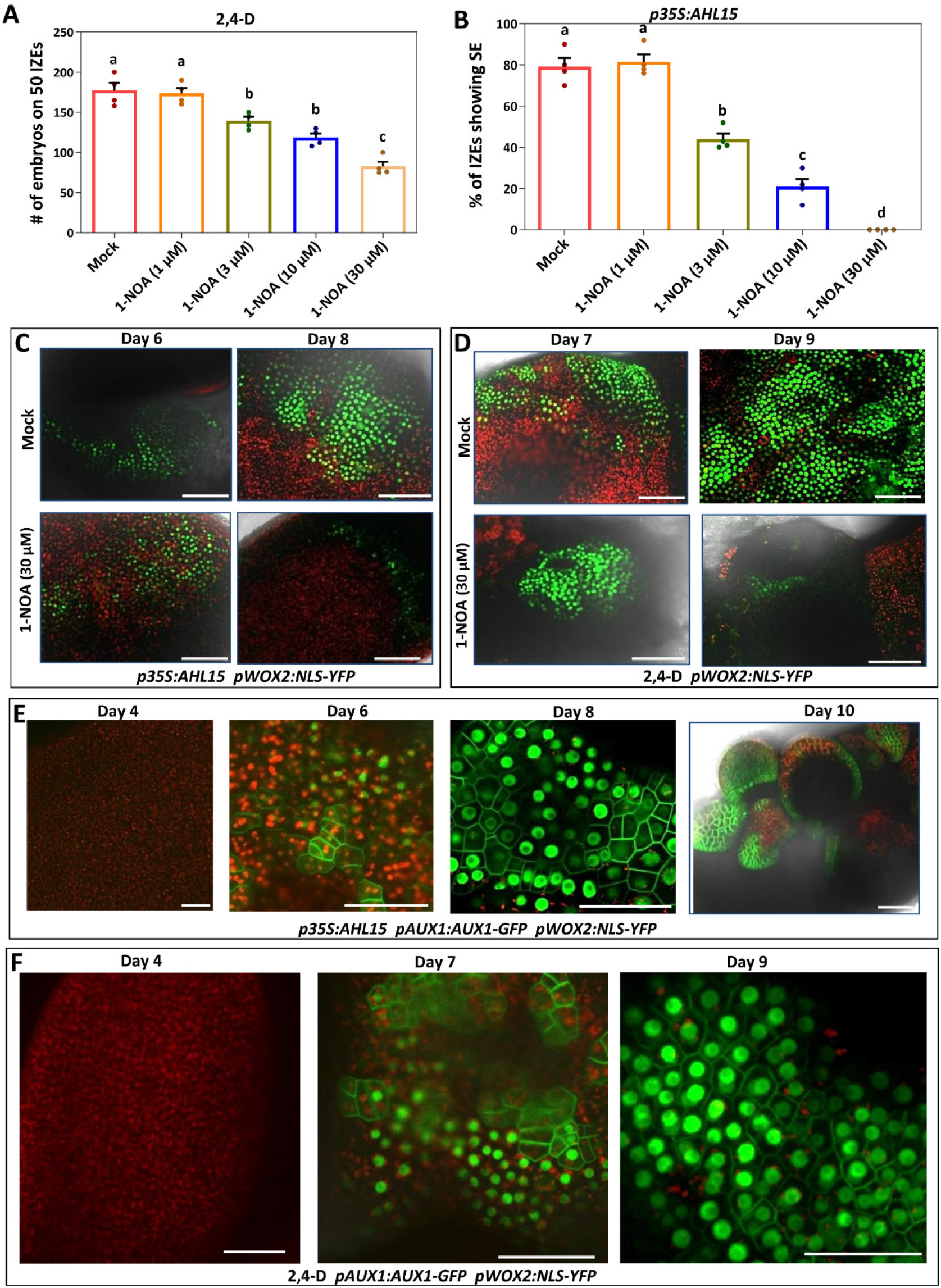
AUX1-mediated auxin influx is required for embryonic cell identity maintenance during SE. (**A**) Number of somatic embryos per 50 wild-type IZEs that were first grown for two weeks on 2,4-D medium with different concentrations of 1-NOA, and subsequently grown for 1 week on medium without 2,4-D (n=4 biological replicates, with 50 IZEs per replicate) (**B)** Efficiency of embryo induction (% of 50 IZEs forming somatic embryos) on cotyledons of *p35S:AHL15* IZEs on medium with different concentrations of 1-NOA. Dots in **A** and **B** indicate the number or percentage, horizontal lines indicate the mean and error bars indicate s.e.m. and different letters indicate statistically significant differences (P < 0.001) as determined by one-way analysis of variance with Tukey’s honest significant difference post hoc test. (**C**) The expression pattern of *pWOX2:NLS-YFP* in cotyledons *p35S:AHL15* IZEs after six or eight days of culture on medium without (up) or with 30 μM 1-NOA (down). (**D**) The expression pattern of *pWOX2:NLS-YFP* in cotyledons of wild type IZEs after six or nine days of culture on medium with 2,4-D only (up) or with 2,4-D and 30 μM 1-NOA (down). (**E**) The expression patterns of *pAUX1:AUX1-GFP* and *pWOX2:NLS-YFP* in cotyledons of *p35S:AHL15* IZEs after four-, six- or eight days of culture or in globular to heart stage somatic embryos formed after ten days of culture. (**F**) The expression patterns of *pAUX1:AUX1-GFP* and *pWOX2:NLS-YFP* in cotyledons of wild-type IZEs after four, seven or nine days of culture on medium with 2,4-D. Size bars indicate 100 μm.

## Discussion

SE is a unique biological process in which differentiated somatic cells acquire embryo identity and develop into embryos. The mechanisms driving acquisition of embryo cell fate in somatic cells is a fundamental question in plant biology. Although recent work has shown that SE involves a complex signaling network and large-scale transcriptional reprogramming, the molecular mechanisms underlying SE are not well understood. Given that an increase in endogenous auxin levels is an important factor for efficient SE (Ivanova et al., 1994; Michalczuk and Druart, 1999; Jiménez and Bangerth, 2001a; Jiménez and Bangerth, 2001b; Cheng et al., 2016; Márquez-López et al., 2018; Vondrakova et al., 2018; Awada et al., 2019), we investigated when and how endogenous auxin promotes SE using two SE systems, 2,4-D- and transcription factor (AHL15)-induced SE on Arabidopsis IZE explants.

### Auxin biosynthesis is required to maintain embryo identity during SE

*De novo* IAA biosynthesis in plant tissues has a large influence on plant growth and development and is essential for proper ZE (Robert et al., 2013; Zhao, 2018). Also for 2,4-D-induced SE, a significant increase in endogenous IAA levels has been reported in various plants species for embryogenic explants compared to non-embryogenic explants (Ivanova et al., 1994; Michalczuk and Druart, 1999; Jiménez and Bangerth, 2001a; Jiménez and Bangerth, 2001b; Cheng et al., 2016; Márquez-López et al., 2018; Vondrakova et al., 2018; Awada et al., 2019). This increase in endogenous IAA levels is thought to be required for SE, and to be mediated by upregulation of the IPyA auxin biosynthesis route.

The rate limiting step in the IPyA pathway is catalyzed by the YUC flavin monooxygenases. The Arabidopsis genome encodes 11 *YUC* genes, and based on the up-regulation of *YUC1/2/4/6/10/11* genes in Arabidopsis embryogenic tissue induced by 2,4-D and a reduced somatic embryo induction in *yuc11* single, *yuc2/4* double and *yuc1/4/10* triple mutants (Bai et al., 2013), it has been suggested that the corresponding genes mediate the increase in the IAA levels required for 2,4-D-induced SE (Wójcikowska et al., 2013). In our 2,4-D-induced SE system *YUC4/6/11* were also upregulated together with *YUC7/8/9*, three *YUC* genes that had not been identified in previous publications Application of the IAA biosynthesis inhibitor yuc (Nishimura et al., 2014) significantly reduced the number of 2,4-D-inuced somatic embryos. *YUC6/7/8/9* gene expression was also found to be upregulated in somatic embryo-forming cotyledons of cultured *p35S:AHL15* IZEs, and application of the yuc or kyn IAA biosynthesis inhibitors severely impaired AHL15-induced SE, while exogenous IAA application alleviated the repression of SE caused by yuc and kyn. Based on our data and given the well-established correlation between the expression of *YUC* genes and IAA levels (Kim et al., 2011; Hentrich et al., 2013), we conclude that the elevated *YUC6*/*7*/*8*/*9* expression levels in cotyledons of cultured 2,4-D-treated or *35S:AHL15* IZEs results in increased IAA levels, which are crucial for the development of somatic embryos on these tissues.

Recently, we showed that up-regulation of *AHL* gene expression is required for 2,4-D-induced SE (Karami et al., 2021). Here we show that the expression of the *YUC6/7/8/9* genes is not up-regulated in response to 2,4-D in the *ahl* loss-of-function mutant background. Therefore, *AHL* genes probably act downstream of 2,4-D and upstream of *YUC*-mediated auxin biosynthesis. Although expression of the auxin response *pDR5:GFP* reporter is clearly reduced in *ahl* loss-of-function mutant zygotic embryos (Supplemental Figure 4B), it remains to be determined whether *AHL* genes also have a role in triggering auxin biosynthesis in zygotic embryos.

Our results show that *pWOX2:NLS-YFP* expression marks three different stages of SE in IZE cotyledon tissues: i) the acquisition of embryonic competence marked by low *pWOX2:NLS-YFP* expression, ii) the formation of somatic proembryos consisting of embryonic stem cells showing high *pWOX2:NLS-YFP* expression, and iii) the development of these proembryos into globular and heart shaped embryos, coinciding with loss of *pWOX2:NLS-YFP* expression (Figure 7; Supplemental Figure 1C). By tracking the activity of this reporter in 2,4-D treated or *p35S:AHL15* cotyledon cells, we showed that induction of embryo identity in cotyledon cells does not require auxin biosynthesis, as it occurs in the presence of the yuc inhibitor, but that under these conditions embryo identity is not maintained resulting in rapid conversion to non-embryonic cells. Therefore, we propose that acquisition of embryonic identity in these cotyledons does not require an increase in IAA levels, but that the maintenance of embryonic identity and progression of embryogenesis requires elevated IAA levels. This is in line with the critical role of auxin biosynthesis in the first steps of zygotic embryo patterning and development (Cheng et al., 2006; Robert et al., 2013).

**Figure 7.**
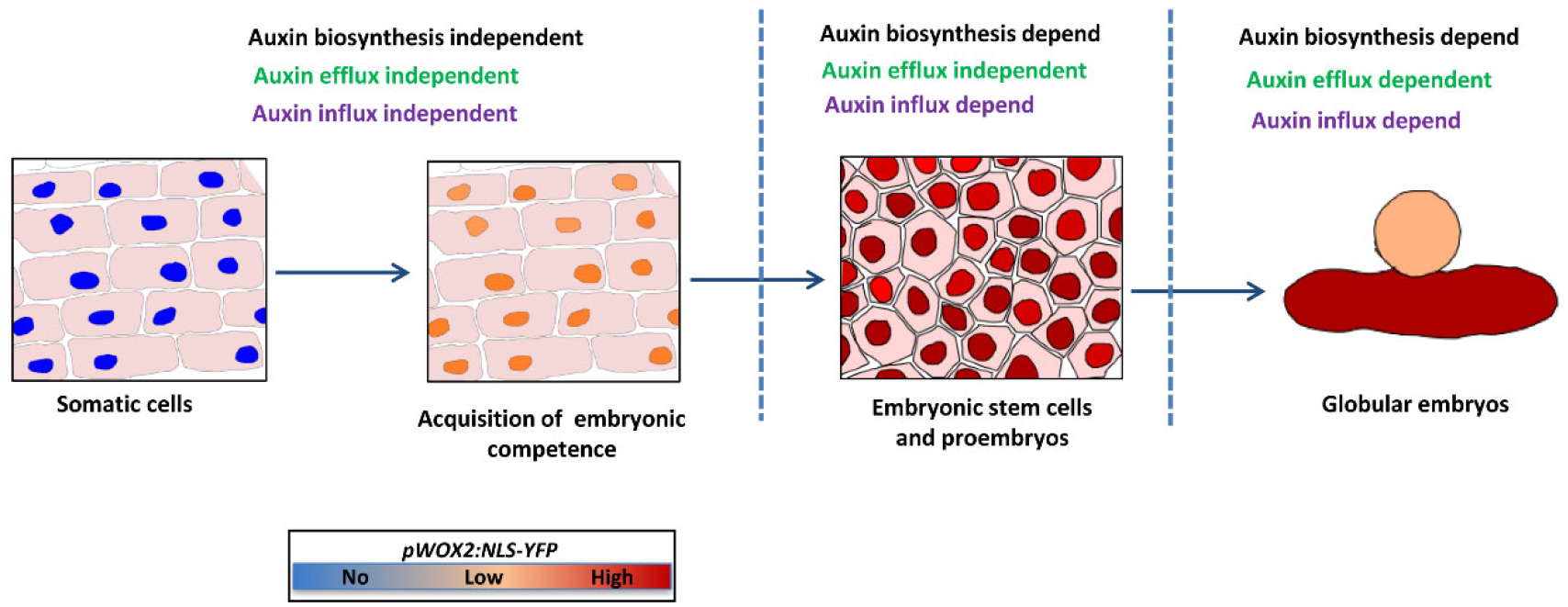
Model for the role of endogenous auxin during different stages of SE. The schematic diagram presents four distinct developmental stages of somatic embryo induction on cotyledons of Arabidopsis IZEs, as distinguished by the *pWOX2:NLS-YFP* embryonic cell identity reporter. The reporter is not expressed in somatic cells. Low expression marks the acquisition of embryogenic competence, which occurs independent of endogenous auxin biosynthesis or transport. The occurrence and maintenance of embryonic stem cells that form proembryos coincides with an increase in *pWOX2:NLS-YFP* expression and requires auxin biosynthesis and auxin influx. The subsequent development of these proembryos into globular embryos and further embryo development leads to loss of *pWOX2:NLS-YFP* expression and is dependent on auxin biosynthesis and transport by the auxin efflux and influx machinery.

This conclusion immediately triggers two questions. If endogenous auxin is not involved in the acquisition of embryonic competence, how can this be triggered by the auxin analog 2,4-D? And why is 2,4-D itself incapable of maintaining embryonic cell identity? The answer to the first question might be that acquisition of embryonic competence requires reprogramming by strong chromatin remodelling (Wang et al., 2020; Karami et al., 2021), something that can only be achieved by transcription factor overexpression or by non-physiological auxin levels. For the second question, the low efficiency of polar cell-to-cell transport of 2,4-D compared to IAA and the specific interaction of 2,4-D with the auxin signaling machinery (Ma et al., 2018) might provide possible explanations. It is well documented that high mitotic activity is necessary for the maintenance of human embryonic stem cell identity (Chen et al., 2015). Consistent with the remarkable similarity in the organization and behavior of stem cells between plants and animals (Heidstra and Sabatini, 2014), high mitotic activity could be essential for the maintenance of the plant embryonic stem cells. Elevated levels of IAA in root and shoot meristems are known to play an important role in stem cell maintenance potentially through promoting cell proliferation (Takatsuka and Umeda, 2014). Therefore, we hypothesize that high IAA levels in embryonic cells might be required to promote cell proliferation.

### Auxin influx and efflux are required for maintaining embryonic cell identify and for embryo development

The directional transport of auxin, facilitated by both influx and efflux carriers, generates and maintains auxin gradients in tissues, and is known to play a crucial role in establishment of the embryonic axis and the development of the zygotic embryo (Möller and Weijers, 2009; Adamowski and Friml, 2015). In contrast, the function of auxin efflux and influx in SE remains largely unknown. In this study, we showed that the auxin influx and efflux machinery plays an important role in the maintenance of embryonic cell identity and proper development of SE. By using the auxin efflux inhibitor NPA or the auxin influx inhibitor 1-NOA, and by tracking expression of the *pWOX2:NLS-YFP* reporter, we found that the early steps in SE, including acquisition of embryonic identity and induction embryonic stem cells do not depend on directional auxin transport. We observed that NPA disrupts the transformation of embryogenic cells into differentiated embryos. In contrast to the normal downregulation of *pWOX2:NLS-YFP* after the globular stage of somatic embryo development, *pWOX2:NLS-YFP* activity was maintained in somatic embryos on NPA-containing medium. From this data we conclude that auxin efflux promotes the development embryonic cell clusters to somatic embryos and later regulates cell fate specification and differentiation during further embryo development.

Among the PIN1-type proteins (PIN1/2/3/4/7) that facilitate auxin efflux in Arabidopsis (Adamowski and Friml, 2015), we only detected expression of PIN1 in embryonic cells and later during embryo development. Previous studies have demonstrated that elevated auxin levels activate the expression of PIN1 proteins (Vieten et al., 2005). Therefore, the appearance of PINI in the embryonic cells may be associated with the auxin biosynthesis-facilitated increase in auxin levels in these tissues.

In ZE, the asymmetric localization of PIN1 on the plasma membrane plays an important role in auxin gradient formation, which is instrumental in cell type specification and pattern formation. (Friml, 2010). PIN1 is expressed in the early one-cell to the 16-cell stage zygotic embryos, where it shows apolar localization. At the 32-cell stage, however, it becomes polarly localized in the provascular tissue to generate an auxin maximum that specifies the hypophyseal cell group. Later, in globular-stage embryos, PIN1 is asymmetrically localized at the plasma membrane of the upper apical region, producing auxin maxima that coincide with the formation of cotyledon primordia (Friml et al., 2003). We did not observe clear polar localization of PIN1 in early embryonic cells during SE, whereas its polar localization on the plasma membrane was detected in globular and subsequent embryo stages. This suggests that in early embryonic cells during SE, just like in early stage (one-cell to the 16-cell stage) zygotic embryos ZE, auxin is not polarly transported, but rather evenly distributed over the embryonic cells. The question arises as to whether PIN1 is the only carrier that facilitates auxin efflux during SE? Other auxin transporters, such as the ATP-binding cassette (ABC) auxin efflux transporters (Geisler et al., 2017), might also contribute to auxin distribution during SE.

Of the four AUX/LAXs proteins that facilitate auxin influx (Swarup and Bhosale, 2019), we only detected expression of *AUX1* during SE. Therefore, we suggest that AUX1 mediates auxin influx during SE. Co-expression of *AUX1* and *pWOX2:NLS-YFP* in embryonic cells suggests that *AUX1* and *PIN1* co-balance auxin influx and efflux in embryonic cells. Unlike NPA treatment, we found that 1-NOA treatment rapidly converted embryonic cells to non-embryonic cells. It seems that auxin influx plays a crucial role in the maintenance of embryonic cell identity. We suggest that conversion of embryonic cells into non-embryonic cells after 1-NOA treatment is related to the reduction of auxin levels in embryonic cells. We hypothesize that PIN1 or other auxin efflux carriers transport auxin to extracellular space, whereas AUX1 prevents auxin leakage by transporting extracellular auxin back to the cytoplasm. This cooperation between auxin influx and efflux in embryonic cells establishes a balance in auxin level in embryonic cells, leading to maintenance embryonic cell identity.

## Conclusions

Taken together, our findings uncover the importance of the endogenous auxin during distinct developmental stages of SE. We show that the acquisition of embryogenic competency and the induction of embryonic stem cells proceed independently of an increase in auxin biosynthesis, or of the auxin efflux and influx machinery. By contrast, an increase in auxin biosynthesis and auxin efflux is essential for the maintenance of embryonic cell identity (Figure 7). Development of embryonic cells into proembryos and the subsequent embryo development also requires an increase in auxin levels together with the auxin efflux and influx machinery (Figure 7). These findings can be used for the optimization of regeneration capacity via SE and for understanding the role of auxin signalling in the regulation of zygotic embryo patterning.

## Materials and methods

### Plant material and growth conditions

All *Arabidopsis thaliana* lines used in this study were in the Columbia (Col-o) background. The transgenic lines *p35S:AHL15, ahl15/+ pAHL15:AHL15-GUS* (Karami et al., 2021), *pDR5:GFP* (Ottenschläger et al., 2003), *pWOX2:NLS-YFP* (Breuninger et al., 2008), *pYUC1:NLS-3xGFP*, *pYUC2:GFP-GUS*, *pYUC4:NLS-3xGFP*, *pYUC5:GFP-GUS*, *pYUC6:GFP-GUS*, *pYUC7:GFP-GUS*, *pYUC8:GFP-GUS*, *pYUC9:GFP-GUS*, *pYUC10:GFP-GUS*, *pYUC11:GFP-GUS* (Robert et al., 2013), *pPIN1:PIN1-YFP* (Benkova et al., 2003) and *pAUX1:AUX1-YFP* (Swarup et al., 2005) have been described previously. Seeds were sterilized in 10 % (v/v) sodium hypochlorite for 12 minutes and then washed four times in sterile water. Sterilized seeds were plated on half MS medium (Murashige and Skoog, 1962) containing 1 % (w/v) sucrose and 0.7 % agar. Seedlings, plants, and explants were grown at 21°C, 70% relative humidity and 16 hours photoperiod.

### Somatic embryogenesis

For the isolation of IZEs at the bent cotyledon stage of development, siliques were harvested 10-12 days after pollination, sterilized in 10 % (v/v) sodium hypochlorite for 7 minutes and then washed four times in sterile water. IZEs were dissected from the siliques inside a laminar flow cabinet (Gaj, 2001). In the AHL15-induced SE system, p35S*:AHL15* IZEs were cultured on solid B5 (Gamborg et al., 1968) supplemented with 2 % (w/v) sucrose and 0.7 % agar (Sigma) for 2 weeks at 21°C, 70% relative humidity and 16 hours photoperiod. Two weeks after culture, the efficiency of SE induction was scored under a stereomicroscope as the percentage of 50 *p35S:AHL15* IZE explants per plate producing somatic embryos. Four plates were scored for each experiment. In the 2,4-D-induced SE system, wild-type IZEs were cultured on solid B5 medium supplemented with 4.5 μM 2,4-D, 2 % (w/v) sucrose and 0.7 % agar (Sigma) for 2 weeks. Subsequently, the embryonic structures were allowed to develop further by transferring the explants to half MS medium with 1 % (w/v) sucrose and 0.7 % agar (Sigma) without 2,4-D. One week after subculture, the capacity to induce SE was scored under a stereomicroscope as the number of somatic embryos produced from 50 IZEs per plate. Four plates were scored for each experiment.

### GUS Staining

Histochemical staining of transgenic lines expressing the β-glucuronidase (GUS) reporter for GUS activity was performed as described previously (Anandalakshmi et al., 1998) for 4 hours at 37 °C, followed by rehydration in a graded ethanol series (75, 50, and 25 %) for 10 minutes each.

### Microscopy

GUS-stained tissues and cultured IZE explants were observed and photographed using a LEICA MZ12 microscopy (Switzerland) equipped with a LEICA DC500 camera.

Confocal Laser Scanning Microscopy (CSLM) was performed with a ZEISS-003-18533. GFP and YFP were detected using a 534 nm laser, a 488 nm LP excitation filter and a 500-525 nm band pass emission filter. Simultaneously, background fluorescence (of e.g. chlorophyll) was captured with a 650nm long pass emission filter. Images were captured with ZEISS ZEN2009 software.

## Acknowledgements

We are grateful to Helene Robert for providing the different *YUC* promoter reporter lines and to Thomas Laux for providing the *pWOX2:NLS-YFP* reporter. We thank Gerda Lamers for help with microscopy and Ward de Winter, Jan Vink and Mariel Lavrijsen for their technical support.

